# Fluorescence Lifetime Imaging Microscopy (FLIM) reveals the diversity and complexities of an open ocean photosymbiosis

**DOI:** 10.64898/2026.09.16.750219

**Authors:** Nicole L Coots, Leocadio Blanco-Bercial, Iliana H. Cutten, Cove Beaven, Patrick J Keeling

**Affiliations:** Department of Botany, University of British Columbia, 2212 Main Mall, Vancouver, BC, V6T 1Z4, Canada; Bermuda Institute of Ocean Sciences, Arizona State University, 17 Biological Station, St. George’s, GE 01, Bermuda

## Abstract

The Retaria are a marine protist lineage that host photosynthetic endosymbionts, where these symbioses play an important role in open ocean food web dynamics. Despite the importance of these relationships, our knowledge about planktonic Retaria photosymbiont diversity remains limited due to the present inability to cultivate these lineages, requiring direct, culture-free observations. Fluorescence in situ Hybridization (FISH) is a common tool that could be used to identify Retarian photosymbionts in nature, but it is challenging to interpret these data due to the substantial autofluorescence of algal plastids. Here, we applied FISH and Fluorescence Lifetime Imaging Microscopy (FLIM) to assess the prevalence of common algal photosymbiont lineages, as this method has the potential to disentangle signal from autofluorescence, even at very similar wavelengths. We applied retarian photosymbiont FISH probes to environmental plankton samples and analyzed the fluorescence lifetime of 92 photosymbionts across 71 hosts. We show that the retarian photosymbiosis is far more complex than originally described, where Retaria commonly host multiple algal lineages per cell and host photosymbionts beyond the most common lineages. These results demonstrate that FLIM is underused in the study of host-microbe associations, with significant potential to distinguish chlorophyll autofluorescence from FISH probe signal in environmental samples.

## Introduction

Protists are present in all major ecosystems, and there is growing evidence that their activities and functions in those ecosystems are frequently impacted by symbiotic associations with other microbial partners. However, most of these symbioses remain uncultivable, so we have few tools to advance our understanding of their interactions. Indeed, many of these systems have only been observed in passing, such that the symbiotic partners have seldom even been identified at the molecular level.

The Retaria are one such protist lineage, where we know living in symbiosis is very common, but for which little information exists about the diversity and mechanistics of these relationships. The Retaria (Rhizaria) are marine microbial eukaryotes classically divided into two subgroups: Foraminifera and Radiolaria. They are found throughout the global ocean, and often in high abundance, such that Retaria and their close relatives account for approximately 5% of the plankton biomass in the top 200 m of the ocean (Biard et al. 2016). This substantial abundance has important implications for marine ecosystem dynamics, where their abundance is correlated with carbon export through the biological carbon pump (Guidi et al. 2016), and their silica skeletons represent up to 19% of the standing silica stock in the ocean (Llopis Monferrer et al. 2020).

Beyond the biogeochemical importance of the group, the Retaria are also noteworthy for their ability to capture and retain algal symbionts via phagocytosis with their large, dense pseudopodial networks. These photosymbioses significantly contribute to net primary production in the open ocean, such that the rate of primary production occurring through retarian photosymbioses is orders of magnitude greater than that of the surrounding water (Caron et al. 1995). This fuels nutrient-deplete, open ocean ecosystems, where the paucity of nitrogen and phosphorous limit phototrophic growth outside of endosymbiosis (Marañón et al. 2003; Duarte et al. 2013).

While the ecological implications of the relationship are clear, our understanding about the mechanisms and interactions that underpin the retarian photosymbiosis is still missing fundamental details. For example, constituents of the retarian holobiont are often inferred only from light microscopy, and single-cell metabarcoding has only been applied in the last five years for some retarian subgroups (Greco et al. 2021; Nakamura et al. 2023; Coots et al. 2025). Given the significant limitations of interpreting light microscopy and the challenges of applying single-cell metabarcode data at a large scale, we sought to refine methods to assess the frequency and diversity of photosymbiosis in Retaria using Fluorescence In Situ Hybridization (FISH).

FISH is a broadly applicable and cost-effective method to identify the partners in microbial interactions; however, the natural emission of light by biological structures (autofluorescence) can complicate the interpretation of fluorescence micrographs outside of samples with known, controlled diversity. Autofluorescence can come from a wide variety of sources, including chlorophyll and lignin in photoautotrophs, or chitin in diverse insect or crustacean cuticles and fungi, among many other sources (Maclsaac and Stockner 1993; Haug et al. 2011; Donaldson and Radotic 2013; Glenn et al. 2013). Autofluorescence can also be derived from methodological sources, such as prolonged sample fixation with aldehydes (Eastham 1968; Weinberg 1989; Staughton et al. 2001). While methods exist to limit or remove some sources of autofluorescence, many sources are unknowable due to the sheer diversity of causes in environmental samples, such that methods to remediate autofluorescence continuously fail (Staughton et al. 2001). These complications necessitate a new approach to disentangle endogenous or exogenous sources of fluorescence from FISH probe signal.

Here, we adapt an approach to describing the retarian photosymbiosis that distinguishes between fluorescence signals using Fluorescence Lifetime Imaging Microscopy (FLIM). FLIM measures the amount of time that a fluorescent molecule spends in an excited state after excitation before emitting a photon of light and returning to a ground state (Figure 1). Repeated laser excitation at stable pulse rates allows us to measure the average amount of time, or the fluorescence lifetime, that the photon stays in the excited state. Broad-spectrum autofluorescence can complicate FISH microscopy because it is difficult to distinguish this “noise” from the “signal” emitted by the probe, but with FLIM we can exploit the fact that different sources of fluorescence, even at highly similar wavelengths, yield different fluorescence lifetimes (Figure 1B-C), and therefore can distinguish fluorescence originating from the target probe from autofluorescence.

**Figure 1.**
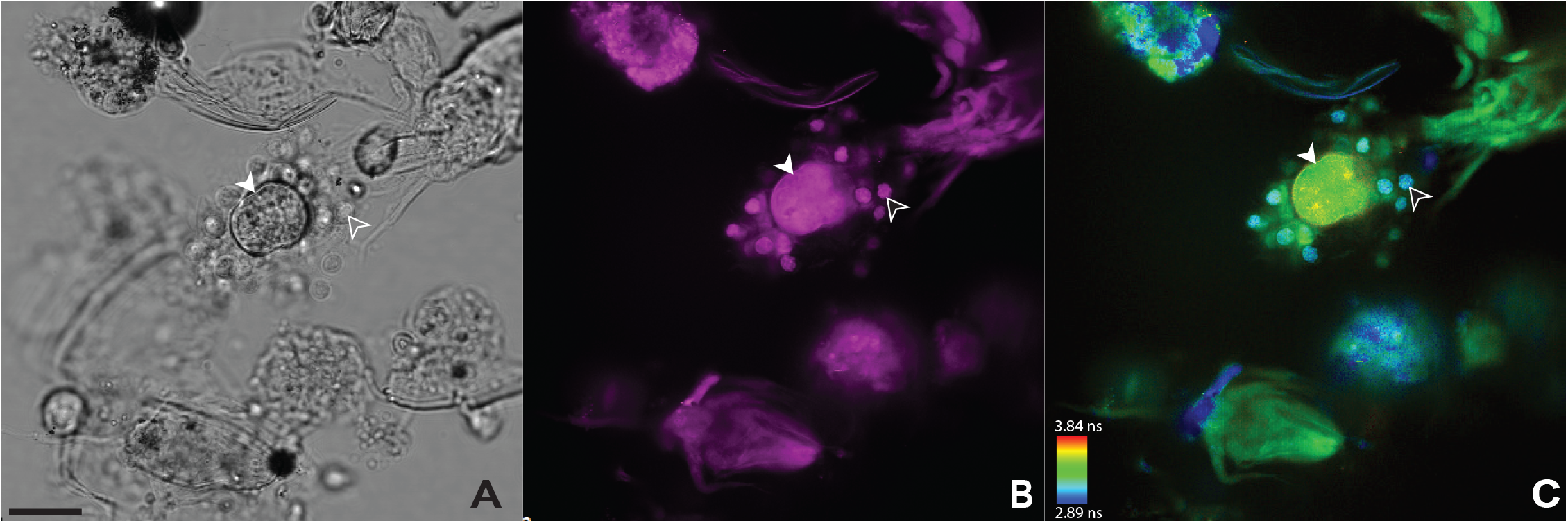
Light and fluorescence micrographs of a fixed, environmental plankton sample collected from the Sargasso Sea. Filled in arrows point to the central capsule of a radiolarian while hollow arrows point to one of the many algal photosymbionts. Scale bar = 50 µm. **A.** Light micrograph taken with brightfield illumination. **B.** Fluorescence micrograph of the far-red emission spectrum of autofluorescence present in the environmental sample. **C.** Heatmap of the fluorescence lifetimes of the sample, demonstrating the variation of different sources of fluorescence, despite visually equivalent wavelengths of fluorescence. Rainbow legend indicates the range of fluorescence lifetimes present in the image collected.

While FLIM is not a new method, it is rarely applied to biological samples, and has not been used with our specific methodology to date. We applied FISH probes targeting common algal lineages to environmental plankton samples, and counted the frequency for which these lineages are intracellular photosymbionts within planktonic Retaria. FLIM measurements were collected for 92 symbiont cells in 71 retarian hosts to validate the source of fluorescence, revealing that planktonic retarians often associate with more than one algal lineage per cell, and that they associate with a much greater diversity of symbionts than previously expected.

### Experimental Procedures

#### Sample collection

Environmental plankton samples were collected by dropping a plankton net (50 cm diameter, 35 µm mesh size) to the approximate depth of the Deep Chlorophyll Maximum (DCM) and towing the net vertically aboard the *R/V Atlantic Explorer* during the monthly Bermuda Atlantic Time-series Study (BATS) cruises (Supplementary Table 2). Plankton samples were concentrated with gravity filtration through a sieve of the same size as the plankton net. Concentrated plankton samples were fixed in 4% neutral buffered formalin with added 250 µM SrCl_2._ The duration of fixation varied from 4 hours to overnight due to the limitations of collecting and fixing samples at sea. After the plankton samples were returned to land, they were washed thrice by allowing the plankton to settle to the bottom of the tube over the course of approximately 1-2 hours, after which point the supernatant was replaced with 0.2 µm-filtered seawater. Fixed, washed samples were stored at 4°C until further processing.

We also fixed stably-growing lab cultures of the dinoflagellate *Oxyrrhis* and its chlorophyte prey, *Dunaliella*. For this, we fixed a small aliquot of the culture with 4% neutral buffered formalin for 4 hours. The sample was then briefly centrifuged to create a soft pellet of the dinoflagellates and the chlorophytes so that the supernatant could be replaced with fresh, 0.5 µm-filtered seawater, which we did thrice. Fixed and washed cultures were stored at 4°C before FISH.

#### Fluorescence in situ Hybridization (FISH)

Before FISH, we first confirmed that all probes that were to be used concurrently would have no competition in terms of the physical location for which they hybridized to the 18S rRNA gene by analyzing their binding locations in Geneious v.2025.1.1. We also confirmed that they all could be used effectively with the same FISH parameters (i.e. formamide content, hybridization temperature, and hybridization duration) and possessed spectrally distinct fluorophores so that little to no fluorophore crosstalk would occur during imaging.

Plankton samples were dried onto Histogrip-coated #1.5 coverslips by incubating the sample on the coverslip in a 46°C incubator for several hours, or until fully dried. Samples were then further dehydrated in a 20 µm HEPES-buffered ethanol series, where the coverslip was incubated in 50%, 75%, and 95% ethanol (twice per concentration) for 3 minutes each. The coverslips were then incubated in the dark in a humidity chamber with a 50 ng/µl probe solution on top of the coverslip (See Supplementary Table 3 for exact reagent solution concentrations). Hybridizations took place for 5 hours at 46°C. After FISH, coverslips were washed in a wash buffer for 30 minutes at 46°C (Supplementary Table 3), briefly submerged in milliQ water at room temperature, then mounted to microscope slides with CitiFluor AF1 mountant. Coverslips were sealed to the microscope slides with clear nail polish.

Three variations of FISH were performed to address different questions. First, *Oxyrrhis* and *Dunaliella* cultures were analyzed as a baseline to validate our method for distinguishing lifetimes. Samples were hybridized concurrently with a general dinoflagellate FISH probe (DINO) conjugated with a FITC fluorophore, a general chlorophyte FISH probe (CHLO) conjugated with a Cy5 fluorophore, and a general eukaryote FISH probe (gen. euk.) conjugated with a Cy3 fluorophore. Coverslips were washed for 30 minutes in wash buffer at 46°C after FISH, then briefly rinsed in milliQ water, and mounted to a microscope slide as described above. Second, we applied the same three FISH probes to marine environmental samples. No DAPI was used for the first two setups. For the third setup, we designed a FISH probe specific to *B. nutricula* using the software platform Oligo-N-design (Sandin et al. 2026). This FISH probe (BNUT) was conjugated with a Cy5 fluorophore and used concurrently with a general eukaryote FISH probe conjugated with a Cy3 fluorophore. Samples in the third setup were stained with DAPI after the milliQ water rinse by placing 300 µl of 100 µg/ml DAPI onto the sample on the coverslip and incubating at room temperature for 30 minutes. Coverslips were then briefly submerged in milliQ water again and mounted to slides.

For every FISH experiment performed, a control coverslip with fixed, washed plankton samples or cultures dried onto it underwent the same protocol for each reaction with all of the same solutions, except without FISH probes. All samples were imaged within 24 hours of mounting.

#### Imaging and Fluorescence Microscopy

Brightfield and fluorescence imaging took place using a Leica Stellaris 8 FALCON confocal microscope with two multiphoton lasers – one set to 780 nm, and one set to 1045 nm. Three HyD detectors were used to collect fluorescence emissions from each probe, where probes with the most similar fluorescence spectra were imaged in two different channels sequentially to prevent fluorescence crosstalk. When two probes were used at once, emission spectra of the two probes were confirmed to be on opposite sides of the wavelength spectrum of visible and UV light to prevent residual biases in fluorescence categorization. Image acquisition parameters including laser intensity and smart gain were first adjusted to optimize signal-to-noise ratio, then kept constant, until lifetime imaging.

Fluorescence Lifetime Imaging Microscopy (FLIM) was used to distinguish between symbiont autofluorescence and FISH probe signal. For this, plankton samples were scanned for Retaria using brightfield microscopy, and each symbiotic retarian present was then imaged with each fluorescence spectrum. The fluorescence lifetime for retarians possessing brightly fluorescing symbionts in one of the fluorophore emission spectra was collected. If a retarian displayed a fluorescence signal in the BNUT, dinoflagellate, or chlorophyte FISH probe emission spectra but not the general eukaryote FISH probe emission spectrum, then the cell and its endobionts were not counted because the general eukaryote FISH probe was used as a control.

Time-Domain FLIM was used to collect FLIM data, where laser pulsing was fixed at 78 MHz. Laser intensity was adjusted to ensure that photon arrival was below or relatively equal to one photon per laser pulse based on a live histogram of photon arrival times, and then the FLIM image and data were collected. After image collection, the average lifetime of all photons present within the boundaries of a retarian symbiont region of interest was measured by applying exponential reconvolution fitting to the fluorescence decay data, which convolves a multi-exponential decay model with the measured instrument response function (IRF) and compares this to raw data. For this, the number of exponents were iteratively adjusted until the measured decay times had the closest fit to theoretical decay times, and until the chi-squared value for curve-fitting was closest to 1.0. This process yields an average and standard deviation lifetime (in nanoseconds) of all pixels in a region of interest. Here, the intensity-weighted average lifetime of all pixels measured within the region of interest (i.e. the symbiont cell) and the associated standard deviation were recorded for every symbiont identified (Supplementary Table 4).

The same protocol was followed to measure the fluorescence lifetime of autofluorescence of retarian symbionts on the control coverslip. Briefly, control coverslips were scanned for symbiotic retarians with brightfield microscopy and, upon finding one, the symbiont autofluorescence lifetime was measured for the emission spectra of FISH probes applied for that specific experimental setup. Symbiont fluorescence on experimental coverslips (i.e. those containing FISH probes) was attributed to the FISH probe hybridization when the standard deviation of the symbiont lifetimes on the experimental coverslip did not overlap with the standard deviation of the symbiont lifetimes on the control coverslip (i.e. the lifetimes of symbiont chlorophyll autofluorescence, with no FISH probes applied). In the case of the experimental setup involving cultured dinoflagellates and chlorophytes, the fluorescence lifetimes of the dinoflagellate and green alga were measured separately and compared to autofluorescence from cells of the same lineages on control coverslips.

## Results

### Validating FLIM-FISH cultured protists

To validate that FLIM-FISH effectively distinguishes probes from autofluorescence, we applied dinoflagellate, chlorophytes, and general eukaryotic FISH probes concurrently to cultured control cells with known identities – the dinoflagellate *Oxyrrhis* and the chlorophyte *Dunaliella*. We compared the fluorescence lifetime of autofluorescence from both lineages on control coverslips to the fluorescence lifetimes observed on experimental coverslips, where the only difference in treatment between the two coverslips was the addition of probes on the experimental coverslip. The average lifetime of the probes was measured within 15 single cells observed on two different dates and compared with the average lifetime of autofluorescence emitted by 15 cells on control coverslips (in each case 8 *Oxyrrhis* cells and 7 *Dunaliella* cells: Supplementary Table 1). Fluorescence lifetimes of cells exposed to lineage-specific probes all differed significantly from the fluorescence lifetimes of autofluorescence on control coverslips (Supplementary Table 1), showing that FLIM can effectively distinguish between different sources of fluorescence.

We next tested the same protocol on retarian cells in environmental samples, using two different sets of probes to identify the symbionts (or lack thereof) from a total of 71 retarians (Figure 2A, C). A large proportion of observed retarians were aposymbiotic: 31 possessed no visible symbionts at any wavelength (Figure 2A, C; 3J-K). Within the 40 retarians for which symbionts were present, we counted a total of 92 symbionts that were labelled with at least one of our FISH probes (Figure 2B, D).

**Figure 2.**
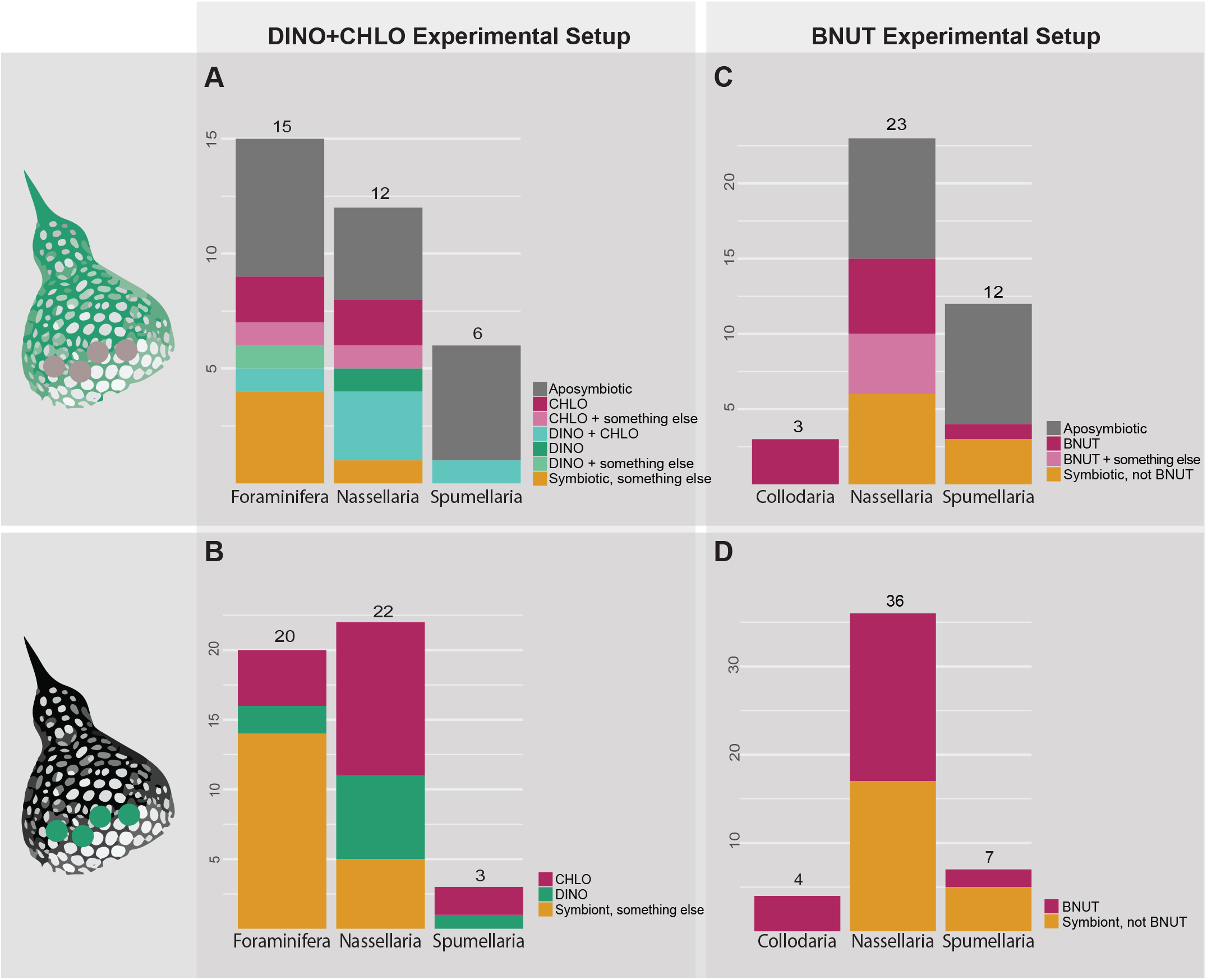
Bar plots of retarian host and symbiont counts performed in this study. Bar plots are grouped by the experimental setup in which counts occurred (columns) and the types of counts that were performed (rows). Numbers at the end of each bar indicate the total number of cells counted for each radiolarian subgroup. Schematic radiolarians demonstrate the type of cell counts recorded in each row, where the focus of the count (either host or symbiont) is highlighted in green. **A.** Bar plot of the total number of retarian cells counted with DINO and CHLO probes added to the sample, and their symbiotic status according to our FLIM analysis, where the total count of radiolarians is subdivided into the subgroups Nassellaria and Spumellaria. **B.** Bar plot of the total number of symbionts counted with DINO and CHLO probes, and the identity of the symbiont according to our FLIM analysis, where the total count of radiolarian symbionts is subdivided into the subgroups Nassellaria and Spumellaria. **C.** Bar plot of the total number of radiolarian cells counted with the BNUT probe added to the sample, and their symbiotic status according to our FLIM analysis, where the total count of radiolarians is subdivided into the radiolarian subgroups Nassellaria, Spumellaria, and Collodaria. **D.** Bar plot of the total number of symbionts counted with the BNUT probes added to the sample, and the identity of the symbiont according to our FLIM analysis, where the total count of symbionts is subdivided based on the radiolarian subgroup from which they were found (Nassellaria, Spumellaria, and Collodaria).

Using the same three probes validated in the *Oxyrrhis-Dunaliella* culture, we measured the average fluorescence lifetime of 45 symbionts across 33 retarian host cells and compared that to the average lifetime of retarian symbionts in control samples (Figure 2A-B). Of the 45 symbiont cells measured, 9 were dinoflagellates and 17 were chlorophytes (Figure 2B). Notably, 19 retarian symbionts hybridized the general eukaryote probe, but the fluorescence corresponding to the wavelengths of the dinoflagellate and chlorophyte fluorophore emission spectra possessed the same fluorescence lifetime as the chlorophyll autofluorescence of retarian symbionts on the control slide. This indicates that 19 eukaryotic symbionts were neither dinoflagellate nor chlorophyte because the corresponding FISH probe did not hybridize.

Retarians occasionally hosted symbionts from diverse algal lineages per host cell. Of the 33 host cells measured, five (four radiolarians and one foraminiferan) harbored both dinoflagellate and chlorophyte symbionts at the same time (Figure 3A-D), whereas three hosts possessed dinoflagellate and/or chlorophyte symbionts in addition to symbionts that visibly hybridized to the general eukaryote probe but not to either lineage-specific probe (Figure 2B). Therefore in sum, 8/33 host cells possessed at least two unrelated algal lineages at once. Of the remaining 25 cells, six hosts had only chlorophyte symbionts, and two had only dinoflagellate symbionts (Figure 2B, 3E-I). Five host cells (four foraminifers and one radiolarian) appeared to exclusively harbor symbionts that hybridized to neither chlorophyte nor dinoflagellate probes (i.e., neither specific probes were detected, but the general eukaryote FISH probe was: Figure 2B).

**Figure 3.**
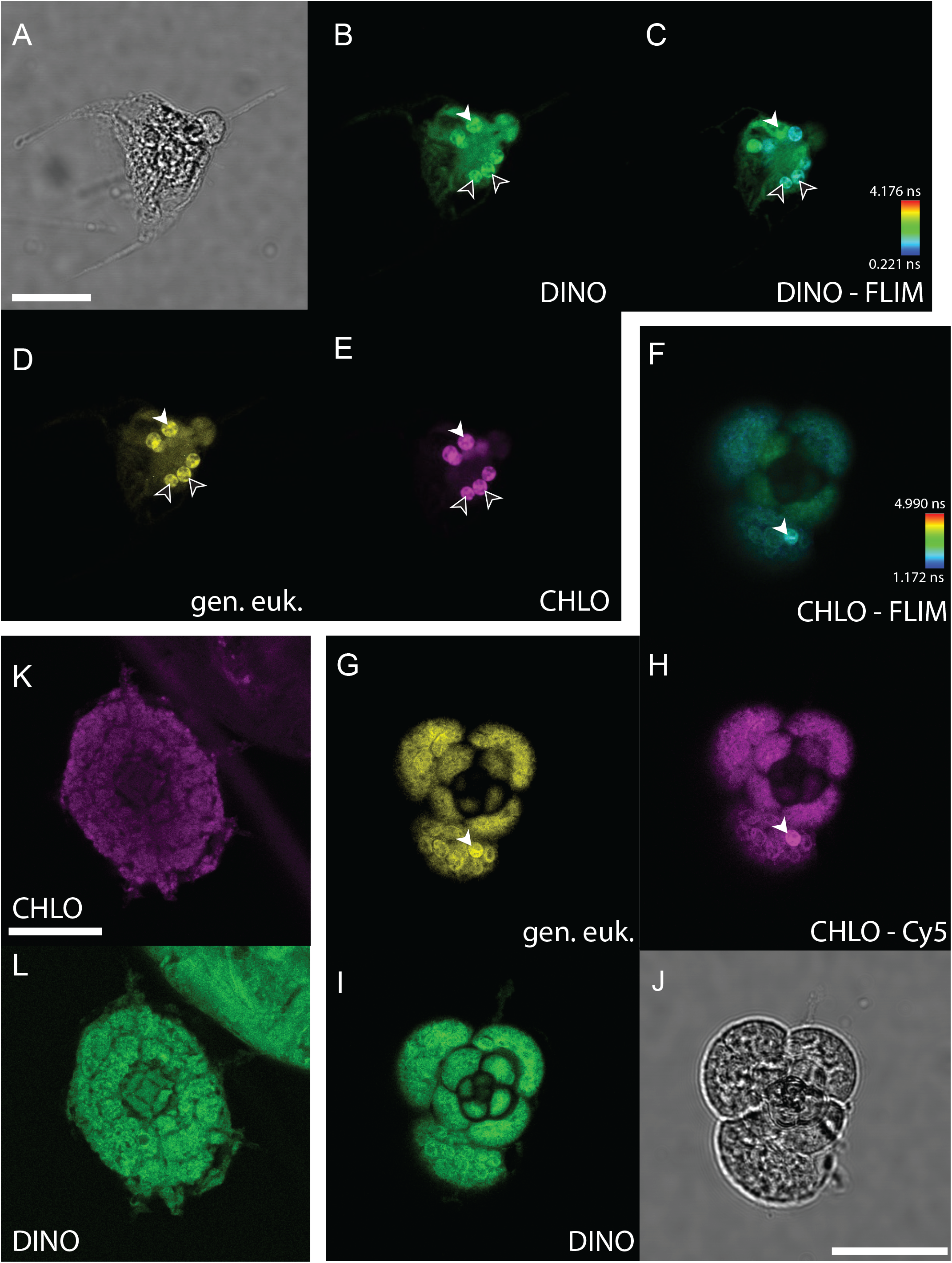
Brightfield and fluorescence micrographs of dinoflagellate-specific (DINO) and chlorophyte-specific (CHLO) FISH probes in Retaria hosts from the Sargasso Sea. Colors for each micrograph are the wavelengths imaged as follows: General eukaryote (gen. euk.) FISH probe conjugated with Cy3 fluorophore (yellow); DINO FISH probe conjugated with FITC fluorophore (green); CHLO FISH probe conjugated with Cy5 fluorophore (far-red/magenta); Heatmap of the fluorescence lifetime of every photon in the either the Cy5 or FITC fluorophore wavelength spectrum in the cell (blue to green spectrum of colors). Scale bars = 50 µm. **A-D.** A radiolarian (subgroup Nassellaria) with both chlorophyte and dinoflagellate symbionts. Filled white arrows indicate a symbiont identified as a chlorophyte according to its fluorescence lifetime, while hollow arrows indicate symbionts identified as chlorophytes according to their fluorescence lifetimes. **C.** Heatmap of the fluorescence lifetime of symbionts visible in the FITC wavelength spectrum of the cell, where symbionts identified as dinoflagellate appear cyan while symbionts identified as chlorophytes appear green. Rainbow legend indicates the range of fluorescence lifetimes present in the image collected. **E-I.** A foraminifer with one chlorophyte symbiont. Arrows indicate a symbiont with a significantly different fluorescence lifetime from the autofluorescence throughout the host cell. **E.** Heatmap of the fluorescence lifetime of emitted photons of light in the Cy5 wavelength spectrum. Rainbow legend indicates the range of fluorescence lifetimes present in the image collected. **J-K.** An aposymbiotic radiolarian (subgroup Spumellaria), with no visible symbionts in the two emission spectra of FISH probes applied in this experimental setup.

### Using FLIM-FISH to distinguish symbionts at the species level

Because we found that the Retaria associate with diverse algal lineages, we next used FLIM-FISH to examine the frequency of the most common radiolarian symbiont, the dinoflagellate *B. nutricula*. Using a *B. nutricula-specific* FISH probe (BNUT) together with a universal eukaryote probe, we observed the frequency of *B. nutricula* in radiolarian hosts. A total of 38 radiolarian cells were observed in this treatment, of which only 13 possessed *B. nutricula* (Figure 2C; 4A-J). Four of the 13 cells with *B. nutricula* also possessed symbionts from other algal lineages. Those symbionts, labeled by the general eukaryotic probe, either lacked the characteristic far-red emission spectrum of the BNUT probe, or exhibited far-red fluorescence with a lifetime that did not differ significantly from that of autofluorescence (Figure 2D; 4A-D).

Nine radiolarians in this experimental setup possessed visible symbionts, but the far-red fluorescence lifetime of all measured symbionts in those cells was not significantly different from that of symbionts in the control coverslip, so they were determined to be non-*B. nutricula* symbionts (Figure 4K-O).

**Figure 4.**
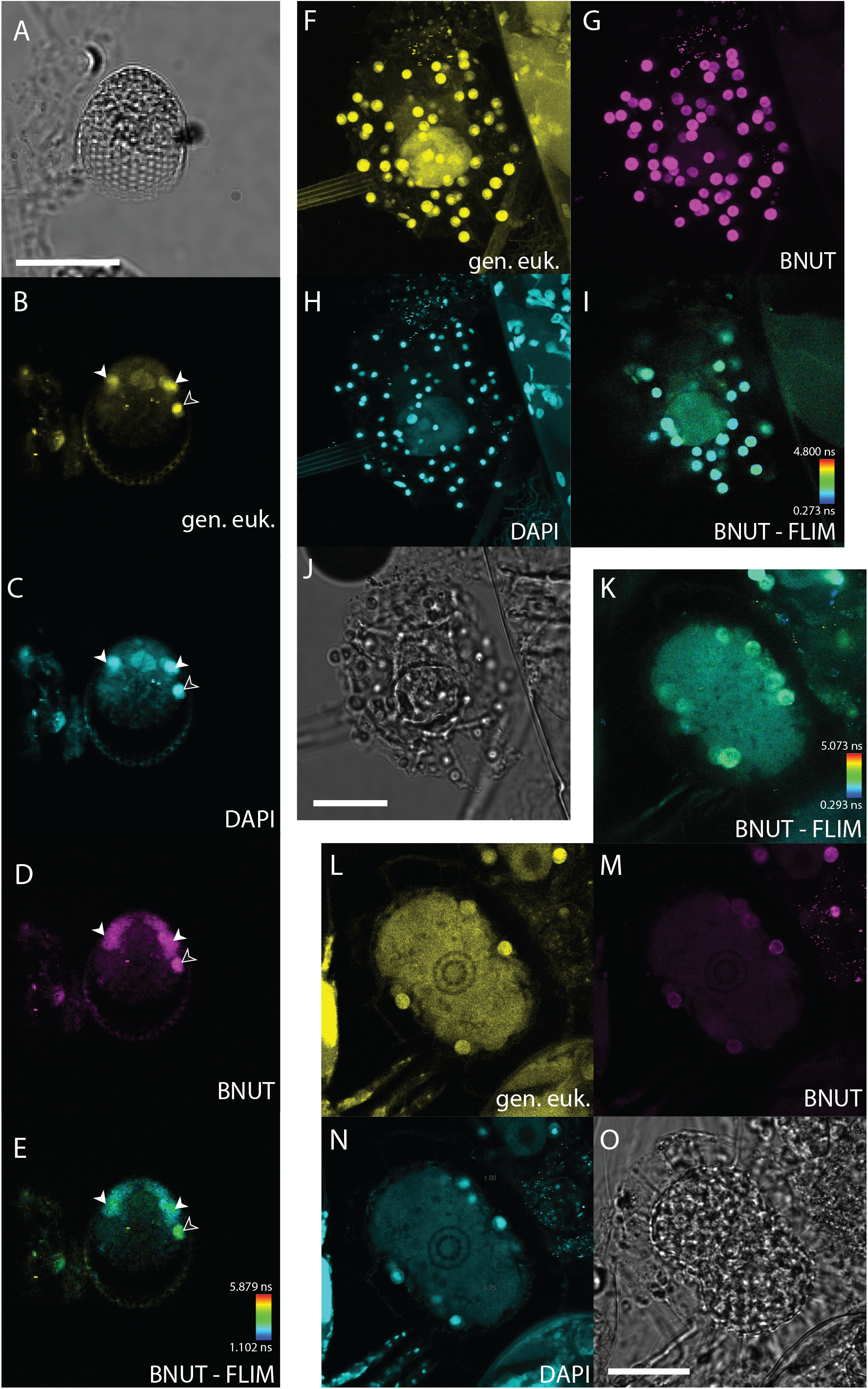
Brightfield and fluorescence micrographs of *B. nutricula*-specific FISH probes in Retaria from the Sargasso Sea. Colors for each micrograph are the wavelengths imaged as follows: General eukaryote (gen. euk.) FISH probe conjugated with Cy3 fluorophore (yellow); DAPI DNA stain (cyan); *B. nutricula-*specific (BNUT) FISH probe conjugated with Cy5 fluorophore (far-red/magenta); Heatmap of the fluorescence lifetime of every photon in the Cy5 fluorophore wavelength spectrum in the cell (blue to green spectrum of colors), where the rainbow legends indicate the range of fluorescence lifetimes present in each of the images in which they were collected. Scale bars = 50 µm. **A-E.** Radiolarian cell (subgroup Nassellaria) with *B. nutricula* symbionts (filled, white arrows) in addition to at least one non-*B. nutricula* symbiont (hollow arrow) according to our analysis of symbiont fluorescence lifetimes. **F-J.** Radiolarian cell (subgroup Nassellaria) with *B. nutricula* symbionts, where all symbionts in which fluorescence lifetime measurements were collected were identified as *B. nutricula*. **K-O.** Radiolarian cell (subgroup Spumellaria) with symbionts according to the gen. euk. FISH probe (**L**) and the DAPI stain (**N)**, and autofluorescence in the Cy5 region (**M**), but no *B. nutricula* symbionts according to the fluorescence lifetime of symbionts present (**K**).

## Discussion

### Retaria are capable of diverse, flexible partnerships

Previous literature suggests that the polycystine Radiolaria predominantly associate with dinoflagellates, followed by chlorophytes in some cases (Gast and Caron 2001; Probert et al. 2014; Yuasa et al. 2016, 2019; Yuasa and Takahashi 2026). Foraminifera are capable of highly diverse photosymbiont associations, where dinoflagellates, chlorophytes, haptophytes, diatoms, and pelagophytes are all reported symbiont lineages (Lee 2006; Pinko et al. 2023; Takagi et al. 2025). For both Radiolaria and Foraminifera, these findings are generally determined by the presence of molecular markers corresponding to those algal lineages (Foster et al. 2006; Bråte et al. 2012; Bourne et al. 2013; Prazeres et al. 2017; Liu et al. 2019). Our findings augment the molecular perspective of Retaria symbiont diversity by adding the fluorescence microscopy perspective. This methodology has revealed that chlorophytes play a bigger role in the Radiolaria symbiosis than previously reported (Figure 2A-B), and that planktonic Foraminifera are more likely to associate with symbiont lineages outside of Chlorophyta and Dinophyceae (Figure 2B).

We also found that the Retaria frequently associate with more than one algal lineage per single host cell (Figure 2, Figure 3A-E, Figure 4A-E). Single-cell metabarcoding of the Retaria often reveal similar findings (Greco et al. 2021; Nakamura et al. 2023; Coots et al. 2025), and indeed, in Coots et al. (2025) the average number of different algal lineages present per radiolarian host was 3.

While photosymbiosis (especially in the Foraminifera) is typically expected to involve a single host lineage and a single algal genus (Garcia-Cuetos et al. 2005; Estrela et al. 2016; Athreya et al. 2025; Takagi et al. 2025), photosymbiont partner flexibility is not unheard of, even in the Foraminifera (Shaked and de Vargas 2006; Schmidt et al. 2018). Furthermore, while the nature of holobiont associations is often drawn into question when studied with metabarcoding or microscopy methods, we suggest that the phytoplankton identified here are more likely to be symbiotic rather than food sources because the Retaria begin digesting prey in their pseudopodial networks, which would affect the intensity of the fluorescence signal of bound FISH probes (Anderson and Be 1976). Here, all counted symbionts reported possessed intense, active chlorophyll based on the chlorophyll autofluorescence, appeared “healthy” in that the cell shape and nuclei were visible and FISH probes bound thoroughly throughout the cell, and were either present inside of the foraminifer skeleton, or they were present in high abundance within the confines of the radiolarian ectoplasm (Figure 3E-I). These traits would not be present in actively digested prey, so our results support that the frequent co-occurrence of algal lineages in Retaria hosts reflects a nonspecific photosymbiotic relationship as opposed to diverse prey sources or eDNA associations.

This flexibility is particularly remarkable given the rich evolutionary history of photosymbiosis in the Retaria, which has played an important role in shaping the lineage for the last 140 mya (Sandin et al. 2025). A flexible, nonspecific strategy for photosymbioses could be beneficial to the hosts – forming low-specificity photosymbioses allows the host to benefit from photosynthesis only when it is advantageous to the current context (Ward 2019) and also allows them to benefit from a wider range of symbionts, if physical access to specific lineages is low. This strategy may be particularly useful in the nutrient-deplete open ocean due to the metabolic plasticity that it provides (Selosse et al. 2017; Edwards 2019; Ward 2019).

### Even the most common radiolarian symbiont is uncommon

Because increasing partner fidelity in photosymbiosis has implications for the degree of host-symbiont interdependency, we next determined the frequency for which the Radiolaria associate with *B. nutricula*. This dinoflagellate species is frequently considered to be the most common photosymbiont lineage to the radiolarian subgroup Polycystinea (Probert et al. 2014; Suzuki and Not 2015; Zhang et al. 2018; Biard 2022), and was the most frequently detected photosymbiont lineage even in molecular surveys that revealed greater algal diversity (Nakamura et al. 2023; Coots et al. 2025).

Using our FLIM-FISH method, we found that *B. nutricula* was only present in 34% of cells identified from Sargasso Sea plankton. This finding further supports the notion that the planktonic retarian photosymbiosis is diverse and lacks host-symbiont species interdependency. Still, it is worth noting that we were only able to count three Collodaria across the year of analyses performed here. Collodaria possess the most specific radiolarian photosymbiosis reported to date, where *B. nutricula* remains the only identified algal lineage in association with Collodaria, and *B. nutricula* was reported to divide *in hospite* within collodarian colonies (Anderson 1982). Indeed, of the three collodarians we found, all three possessed *B. nutricula* exclusively (Figure 2B) and single-cell metabarcoding indicates a similar finding (Coots et al. 2025). Thus, Collodaria have likely evolved a more interdependent relationship with their algal symbiont than the rest of the Radiolaria.

### The application of FLIM as a tool to distinguish between sources of fluorescence

Despite the huge variety of applications of FISH, interpreting fluorescence signal in environmental samples is a notoriously challenging problem to address with traditional FISH methods due to the potential complexity of these systems, coupled with the high frequency of broad-spectrum autofluorescence, especially in phototrophic cells with chlorophyll (Figure 1) (Noonberg et al. 1992; Baschong et al. 2001; Alam et al. 2016). FLIM FISH effectively addresses these challenges and is thus an underutilized tool for the identification of FISH probe signal in cells with autofluorescence. This utility has a wide diversity of applications, including applying FISH probes to environmental samples to search for host-microbe associations, as we have done here.

To date, just one manuscript reports the use of FLIM to study marine host-microbe associations. Deore et al. (2022) used FLIM Phasor plot analyses to distinguish coral autofluorescence from algal symbiont fluorescence. Phasor approaches for FLIM involve the microscope software platform producing a 2D graphical plot that transforms raw fluorescence decay data into Cartesian coordinates (Ranjit et al. 2018). This approach makes the analysis fast, visual, and “math-free” for the user (Digman et al. 2008). However, phasor approaches for FLIM are insufficient for most environmental FISH applications with complex fluorescence profiles because they are unable to provide explicit, independent lifetimes of a single region of interest, such that fine-scale differentiations between autofluorescence and FISH probe signal can become impossible in samples with a large diversity of fluorescing molecules. Because fluorophore lifetimes are sensitive to their environment, the fluorescence lifetime readout can vary by tenths of a nanosecond, making these fine-scale differentiations highly pertinent (Noomnarm and Clegg 2009; Berezin and Achilefu 2010). Additionally, Phasor analyses are sensitive to low signal-to-noise ratios, such that low photon counts per pixel are more likely to affect the final FLIM readout for a cell (Gonzalez Pisfil et al. 2022). This is especially problematic if there is substantial autofluorescence in the rest of the image, as there often is with environmental samples.

For these reasons, we applied a curve-fitting model for distinguishing between the chlorophyll autofluorescence of an alga and a hybridized FISH probe on the same cell. Our chosen method involves no new tools, but instead, a new way of applying known tools to a new purpose. By applying multi-exponential decay curve fitting methods, we were able to determine the precise fluorescence lifetime of every pixel within the confines of the symbiont cell, irrespective of the highly varied sources of fluorescence throughout the environmental plankton sample. By simply comparing this fluorescence lifetime to the equivalent measurements on a control coverslip with no FISH probes, we were able to determine whether the fluorescence signal is new based on the applied probes or whether the fluorescence originated from the sample itself. This method is straightforward to use for addressing the notorious issue of chlorophyll autofluorescence, making FISH on algal cells more accessible.

**Supplementary Table 1.** Fluorescence lifetimes of each measured cell.

## Data Availability

All data collected for this analysis is present in Supplementary Table S1.

## Acknowledgements

This work was supported by a grant from the Gordon and Betty Moore Foundation (https://doi.org/10.37807/GBMF9201) to PJK. We thank the Bermuda Atlantic Time Series (NSF OCE 2241455) for providing monthly fixed environmental plankton samples for this analysis. The authors also acknowledge the UBC Bioimaging Facility, for which this research would not have been possible. LBB acknowledges the funding from NSF OCE 2227766 and the Simons Foundational International’s BIOS-SCOPE program.

